# MCWs (MiCroWire sorter): A new framework for automated and reliable spike sorting in human intracerebral recordings

**DOI:** 10.1101/2025.07.09.663285

**Authors:** Alexander Betancourt, Masoud Khani, Tapasi Brahma, Fernando J. Chaure, Connor Hauder, Sunil Mathew, Ana Sofia Dominguez Zesati, Sean Lew, Kunal Gupta, Hernan G. Rey

## Abstract

Efficient and accurate spike sorting is critical for isolating single neurons from extracellular recordings to distinguish neural activity of interest. However, while the electrodes and acquisition systems for non-human electrophysiology have been enhanced over the past decades to enable higher-yield single-neuron detections, those advances have not been translated into human electrophysiology. Single-wire electrodes are still ubiquitously used, and although acquisition systems have augmented their signal-to-noise ratio over the last 15 years, we are still limited by their low electrode count. Moreover, unlike non-human recordings, human recordings often take place in hospitals where different noise sources and subject breaks can compromise the recording quality during experimental sessions. To bridge this gap, this work presents an automatic, open-source spike sorting pipeline that leverages contemporary computational capabilities and is tailored to single-neuron recordings from humans acquired via microwires. The pipeline is implemented in both MATLAB and Python, ensuring accessibility and compatibility across computational environments. Its modular and comprehensive structure supports customization and even opportunities for new developments as per the requirements of the user and the application. One feature is a data-driven automatic module to remove narrow-band interference, besides electrical line noise, which can be an essential tool while recording in clinical settings, particularly for online processing implementations. Following spike detection, the pipeline implements an artifact rejection module that separates waveforms that are unlikely to be associated with actual spikes. Additionally, we introduce a configurable feature-extraction, clustering, and benchmarking framework that not only allows flexibility in employing user-defined or conventional algorithms, such as wavelet transform with superparamagnetic clustering, but can also evaluate multi-method agreement among the different sorters. The pipeline also utilizes established and novel quality metrics to support semiautomatic curation of isolated clusters. Furthermore, we can integrate the customized pipeline with experimental tasks by removing task-unrelated waveforms (e.g., during a break in a task) and prevent over-clustering with the aid of metrics for comparing response profiles. Thus, the presented pipeline addresses the three-pronged objectives of algorithm-adaptability, rigorous validation, and human single-neuron recording optimization to support clinical and cognitive neuroscience applications.

## 1. Introduction

Invasive monitoring in patients with refractory epilepsy provides a unique opportunity to study the human brain at single-neuron resolution, and over the last 20 years, it has been an important means to study human cognition [1], [2], [3], [4]. These discoveries have demonstrated that individual neurons in the medial temporal lobe encode abstract concepts with remarkable specificity and consistency across different sensory inputs [1], [2]. This would not have been possible without precise spike sorting in human microwire recording to establish accurate response selectivity profiles. In addition, over a decade ago, the use of microwires to analyze activity during seizures started to rise [2], [5], and now it has become even more popular [6], [7], with strong potential to inform targeted therapeutic interventions [6], [8]. The ability to reliably isolate single-neuron activity from human recordings thus represents a critical capability for advancing both fundamental neuroscience and clinical applications [2], [9].

Current animal electrophysiology has achieved remarkable technological sophistication, with modern silicon probes routinely accommodating over 1,000 recording sites per shank and Neuropixels 2.0 incorporating 5,120 sites across four shanks [10]. This high-density electrode technology has driven the development of GPU-accelerated spike sorting algorithms specifically optimized for dense geometries [11], resulting in substantially higher single-neuron yields compared to human recording. Human electrophysiology, by contrast, remains constrained by clinical depth electrodes containing just 8–16 microwires of 40 μm diameter each, fundamentally limiting spatial resolution and discrimination capabilities [2]. Despite improvements in acquisition system signal-to-noise ratios (SNR) over the past 15 years, the low electrode count continues to restrict neuronal yield in human studies. Additionally, clinical recording environments introduce unique challenges, including electromagnetic interference from medical equipment [6], patient care interruptions, and dynamic noise conditions that compromise automated sorting algorithms designed for controlled laboratory settings.

To address these challenges, several spike sorting frameworks have been developed specifically for human microwire recordings, with their efficacy and methodologies documented across various neuroscience publications. **Combinato** [12], [13], an end-to-end solution developed by the Mormann laboratory, has proven effective in large-scale memory and epilepsy studies. Its Python/CPU implementation scales across many cores but lacks GPU acceleration, and the system frequently requires manual intervention to resolve a tendency toward over-clustering [14]. **OSort** [15], developed by the Rutishauser laboratory, uses a semi-supervised approach that excels with variable SNR, but its reliance on significant manual parameter tuning for different recording conditions hinders full automation. The **Wave_Clus 3** [16], [17] framework, developed by Quian Quiroga and Rey, achieved automation through data-driven feature selection, but its current approach, based on non-Gaussian wavelet coefficient distributions, is suboptimal for effective clustering. Differing from these specific algorithms, the **Human Single Unit Pipeline (HSUPipeline)** [18] offers a broader, workflow-oriented solution that standardizes analysis by integrating tools like SpikeInterface [19], [20] and the Neurodata Without Borders (NWB) data format. Its primary limitation is its complete reliance on these external packages. The pipeline’s overall performance and success are therefore dependent on the efficacy and continued maintenance of the underlying third-party sorters it employs.

Despite these advances, existing approaches present significant limitations that compromise their effectiveness in invasive human electrophysiology. Current solutions either lack optimization for sparse microwire arrays, require extensive manual intervention, or fail to robustly handle the dynamic noise conditions typical of hospital settings. No existing framework adequately addresses the combination of algorithm adaptability, rigorous validation, and human-specific optimization required for reliable clinical deployment. The need for customizable, modular solutions that can accommodate diverse methodological approaches while maintaining automated operation in challenging recording environments represents a critical gap in current capabilities. This work addresses these limitations by presenting MCWs (MiCroWire sorter), a comprehensive spike sorting pipeline specifically engineered for human microwire recordings, incorporating data-driven interference removal, sophisticated artifact rejection, and configurable processing frameworks that support both automated operation and user customization.

Our proposed framework is specifically targeted for microwire recordings during Stereoelectroencephalography (sEEG). This is a procedure where depth electrodes are implanted in patients with refractory epilepsy who are undergoing invasive monitoring for clinical purposes (to identify the area of the brain responsible for seizure onset). To accomplish this, commercially available depth electrodes are used, which have iEEG (intracranial electroencephalography) contacts to monitor brain activity across different areas of the brain. By using a modified probe on a given trajectory, we can include 8 high-impedance microwires (and 1 low-impedance microwire that can serve as a local reference), each being 40 μm in diameter, which are trimmed so they protrude 3–5 mm from the tip of the clinical probe. The signals acquired by the wires can be sampled at high rates (30 kHz), carrying sufficient high-frequency activity (> 300 Hz) to enable the study of single-unit activity (SUA). These electrodes are stereotactically implanted in regions of interest, including the hippocampus (body and head), parahippocampal gyrus, fusiform, and amygdala, allowing for precise targeting of deep brain structures typically implicated in memory and perception.

A key advantage of using microwires over Utah Array electrodes is their flexibility and suitability for deep brain recordings. Microwires can be bundled with clinical depth electrodes and advanced into specific deep brain targets with minimal additional tissue disruption, which is not feasible with the rigid, surface-limited Utah Array. This makes microwires the preferred choice for acute, single-unit recordings in deep brain regions in human subjects. However, microwires are mechanically delicate and more susceptible to movement, bending, and electrical noise from the hospital environment or subject activity compared to the sturdier Utah Array. These challenges can result in increased signal variability and artifact contamination, underscoring the necessity for a robust and adaptive spike sorting pipeline, such as the one presented in this manuscript, to reliably isolate and analyze single-neuron activity from microwire recordings in complex clinical settings. Figure 1 shows the sEEG depth microwires, surgically implanted in the left Amygdala and the left mid-Fusiform Gyrus. After surgery, a post-operative CT is co-registered with the preoperative MRI to assess the accuracy of the actual position of the electrodes [2], [21]. The microwire bundles generate an artifact in the CT as they protrude from the tip of the clinical probe.

**Figure 1:**
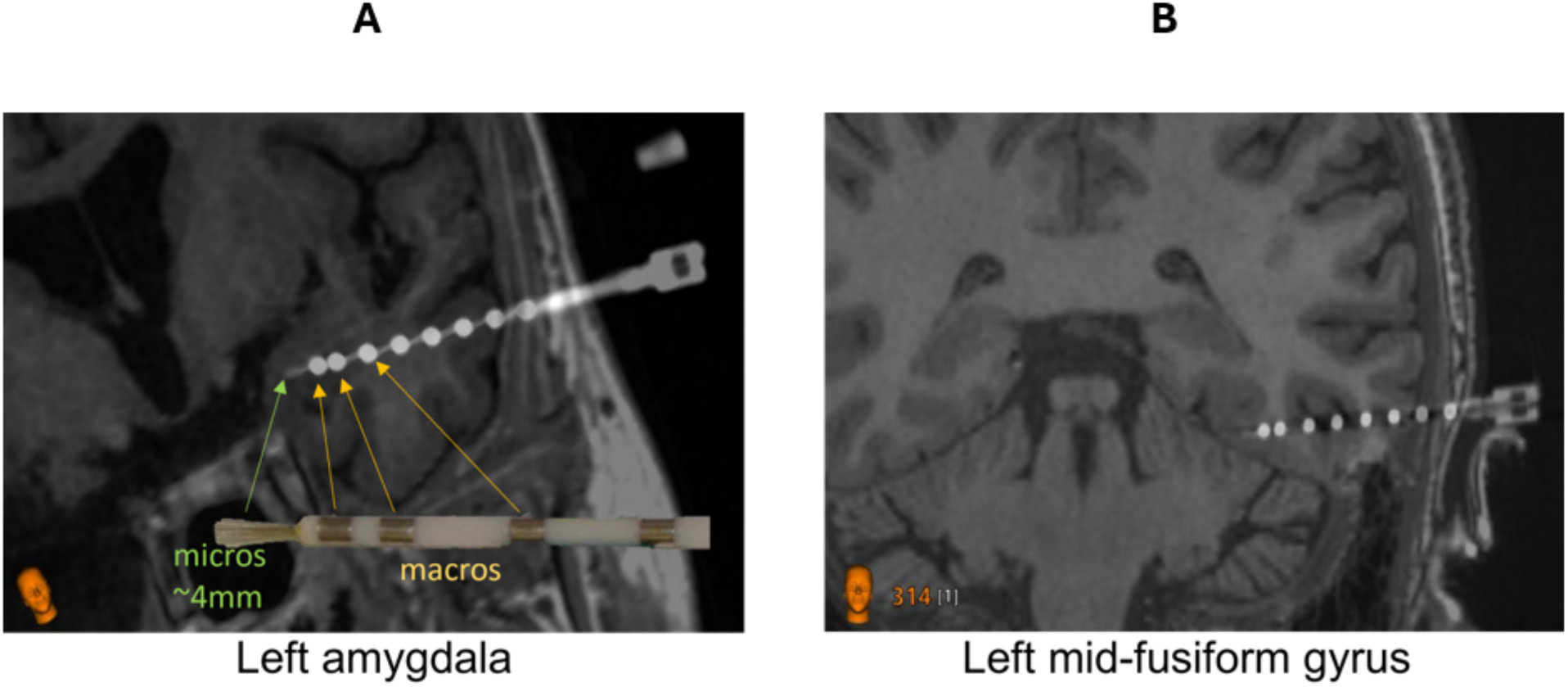
CT-MRI coregistration showing sEEG depth microwires, bundled with clinical depth macro-electrodes, in (A) the left Amygdala and (B) the left mid-Fusiform Gyrus. (A) Yellow and green arrows highlight the clinical macro-electrodes and sEEG depth microwires, respectively, on the CT-MRI coregistered scan, referencing from a photograph of the bundled microwires and macro-electrodes.

## 2. Methodology

The core of our methodology is a novel spike-sorting pipeline meticulously designed and optimized for processing microwire recordings obtained from human subjects in clinical settings. Recognizing the diverse computational preferences within the neuroscience community, the pipeline is offered with a dual implementation in both Python and MATLAB, ensuring broad accessibility and facilitating adoption across different research environments.

A key characteristic of our pipeline is its modular architecture. It is structured as a series of interconnected yet independent processing stages, each dedicated to a specific task in the spike sorting workflow. This modular design philosophy offers significant advantages, including enhanced flexibility for users to customize the pipeline by selecting, modifying, or even replacing individual modules with alternative algorithms or custom-developed tools. Furthermore, it simplifies maintenance, updates, and the integration of future advancements in spike sorting techniques. A detailed flowchart illustrating the sequence and interaction of these modules is presented in Figure 2. The pipeline systematically processes raw electrophysiological data through several key stages: (1) initial data loading and preprocessing, including robust noise reduction; (2) spike detection using adaptive thresholding mechanisms; (3) sophisticated artifact rejection to distinguish neural spikes from non-neural events; (4) feature extraction from detected spike waveforms using a selection of methods; (5) clustering of spikes into putative single-units; (6) comprehensive quality assessment of sorted units using a suite of metrics; and (7) optional integration with experimental tasks to refine sorting, including metrics based on neural response profiles. Subsequent sections will elaborate on the specific algorithms and methodologies employed within each module of this architecture.

**Figure 2:**
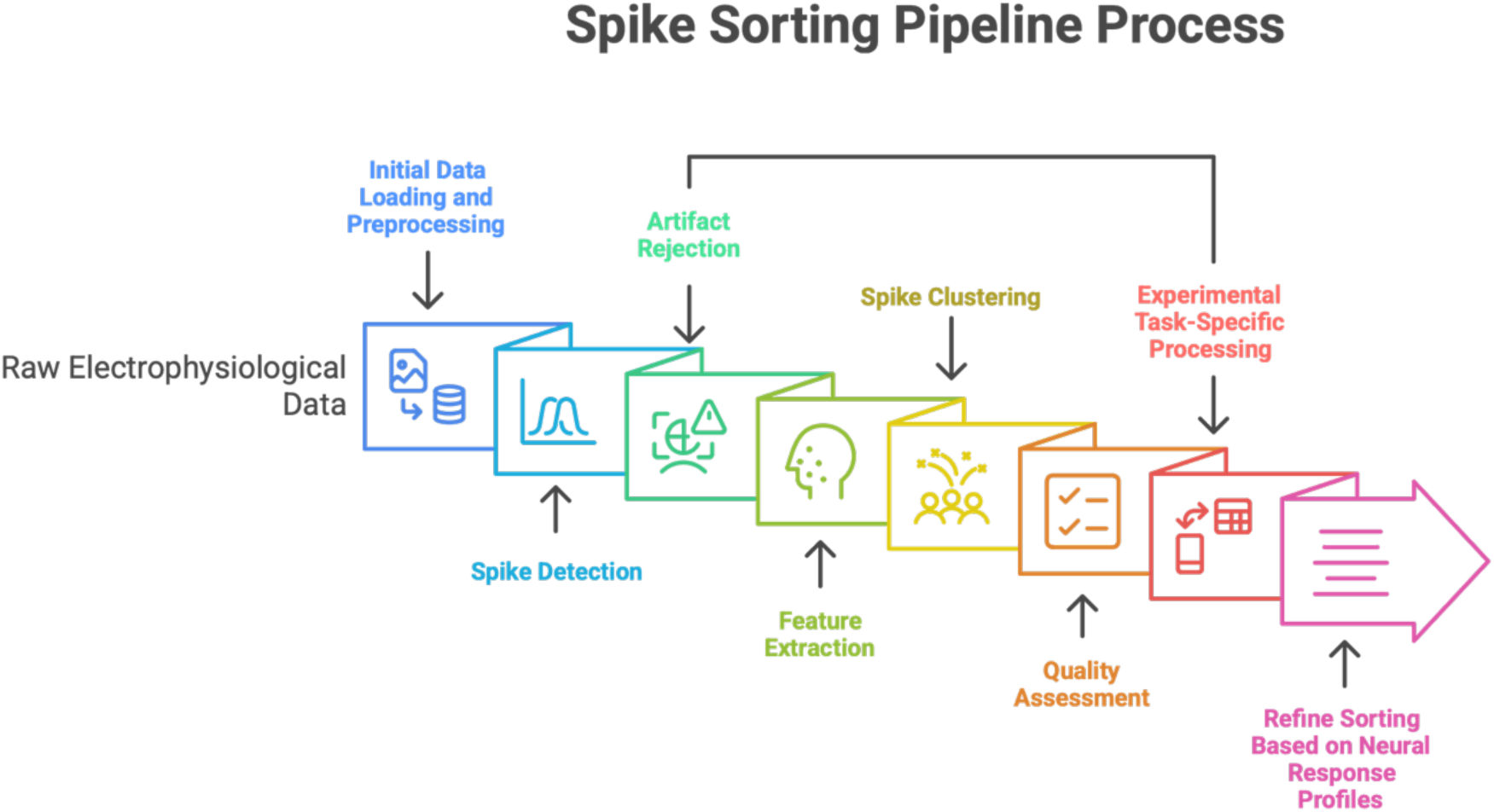
Schematic of the sequential functional modules in the MCWs pipeline: The pipeline processes raw electrophysiological data through seven functional modules: (1) data loading and noise reduction; (2) spike detection via adaptive thresholds; (3) artifact rejection to isolate neural events; (4) feature extraction from spike waveforms; (5) clustering into putative single units; (6) quality assessment of sorted units; and (7) optional task-based integration to refine sorting based on neural responses.

### 2.1 Data-Driven Noise Removal

Our signal processing pipeline begins with the data-driven noise removal module that combines high-resolution power spectrum analysis with adaptive notch filtering to robustly suppress interference while preserving physiological signals. Unlike traditional approaches that rely on assumptions about specific noise frequencies (e.g., line noise and harmonics), our method accounts for the varied potential noise sources in a hospital and automatically detects narrowband noise frequencies in a data-driven way.

The module starts with applying a bandpass filter to the recorded signal to retain the most relevant frequency band for spiking activity between 300 and 3000 Hz [9]. Thereafter, the bandpass filtered signal and the raw signal undergo the Welch method to obtain the power spectral density (PSD) of the signals, applying a Bartlett-Hann window to create segments leading to a resolution of at least 0.5Hz, and averaging the computed periodograms over 200 signal segments (Figure 3A).

**Figure 3:**
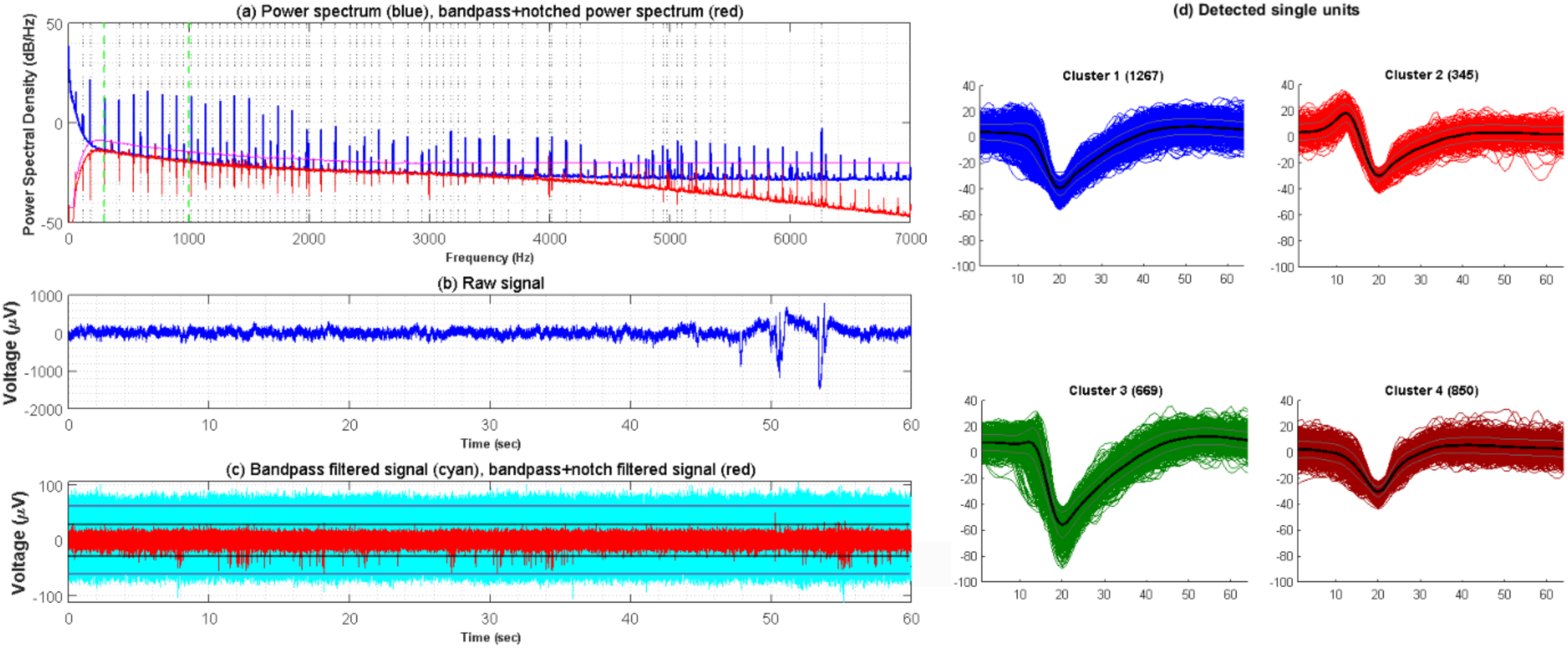
Data-Driven Noise Removal in action: (a) Contrast between the raw signal PSD (blue) and after applying the bandpass and 63 customized notch filters (red). (b) The raw time-series signal before implementing the Data-Driven Noise Removal module. (c) Contrast between the bandpassed time-series signal before (cyan) and after (red) applying the cascade of customized notch filters. Detection thresholds can be seen, highlighting that after removing the narrowband noise, the spikes in the signal (red) can be easily identified (3553 detected waveforms), whereas without the notches, it led to 125268 waveforms detected, nearly all of which were simply due to noise. (d) Detected single-units from the bandpass and notch-filtered signal. The various SUA can be clearly identified after passing the signal through the Data-Driven Noise Removal module.

Next, we apply a moving median filter to the PSD of the bandpass signal, creating a robust and smoothed estimate of the uncontaminated signal that is inherently resistant to outlier values. This median-based approach is particularly valuable as it prevents transient artifacts from disproportionately influencing the baseline while preserving genuine physiological activity. The module then calculates an automated threshold for noise peak detection by adding a fixed margin of 5 dB above this smoothed PSD estimation, eliminating the need for manual noise frequency identification. For each detected noise peak, an adaptive notch filter is designed with its bandwidth tailored to both the width (bandwidth of the noise peak above threshold) and height (relative amplitude to the largest peak) of the interference, ensuring that broader and stronger peaks are more thoroughly suppressed while minimizing unnecessary signal loss for narrow or weak peaks. Finally, the notch bandwidth is constrained between 1 and 3 Hz at lower frequencies and up to 5 Hz at higher frequencies, balancing effective noise rejection with the preservation of critical neural information.

This Data-Driven Noise Removal module assumes the stationary nature of narrowband interference and hence, the noise peaks detected across any segment of the recording can be utilized to design notch filters for the entire recorded signal. Lastly, the parameters of the designed narrowband interference filters are saved to be passed on as inputs to the following functional modules downstream along the pipeline to ensure uniform signal processing by all modules while avoiding any discrepancies. Figure 3 demonstrates the impact of the Data-Driven Noise Removal module in filtering out the narrow-band interference from the recorded signal, which down the line facilitates accurate detection of single-units. Figure 3C highlights the contrast between the signal PSD before and after applying the notch filters, showing that no spikes would have been detected in the bandpassed signal without the notches. In the example presented in the figure below, after removing the narrowband noise, the spikes in the notch-filtered signal (red) can be easily identified (3553 detected waveforms), whereas without the notches, 125268 waveforms were detected, nearly all of which were due to noise. Moreover, Figure 3D shows that good-quality spikes were obtained after the Data-Driven Noise Removal module, with enough information to isolate different putative units.

### 2.2 Spike Detection

Spike detection constitutes a critical initial step in our processing pipeline, fundamentally impacting the quality and reliability of subsequent sorting steps. This module identifies putative action potentials within the bandpass–filtered extracellular recordings by isolating transient voltage deflections that exceed background noise levels.

#### Noise-Floor Estimation

To establish a robust threshold for spike detection, we first estimate the noise floor for each recording channel. Rather than using the standard deviation, which can be substantially distorted by large spikes in the signal, we implement the median absolute deviation (MAD) method [9], [22]:

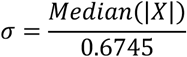

where X represents the bandpass-filtered signal. This approach provides a more accurate estimate of noise levels in the presence of neuronal spikes and transient artifacts, as the median is inherently resistant to outliers compared to mean-based measures. Our pipeline calculates this noise estimate individually for each recording channel to account for electrode-specific characteristics and recording-quality variations commonly encountered in human microwire recordings.

#### Threshold Selection and Adaptation

Based on the robust noise estimate, our pipeline sets amplitude thresholds proportional to the estimated noise level:

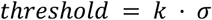

where 𝑘 typically ranges from 4 to 5, allowing users to adjust the sensitivity according to recording conditions and experimental requirements. This parameter balances detection sensitivity and false positive rates, with higher values yielding fewer, but more reliable, spike candidates.

To accommodate non-stationary noise characteristics common in extended clinical recordings, our pipeline implements an adaptive threshold mechanism. The noise floor is re-estimated at regular intervals (by default, every 5 minutes of continuous recording) to track gradual changes in signal quality. This adaptation is particularly valuable in human recordings, where subject movement, environmental factors in hospital settings, and the drift of electrodes can alter background noise levels throughout experimental sessions.

#### Candidate Extraction and Alignment

Once threshold crossings are identified, our pipeline implements a peak search algorithm [22] to locate the precise timing of each putative spike. For each threshold crossing event, we extract a temporal window of approximately 2-3 ms (configurable parameter) centered on the detected peak to capture the complete spike waveform for subsequent analysis.

To enhance the precision of temporal alignment, critical for accurate feature extraction and clustering, we implement sub-sample interpolation using cubic splines. This approach achieves alignment precision of approximately ±0.5 samples, substantially reducing jitter-induced variance in the extracted waveforms. Precise alignment is particularly important for human microwire recordings, where relatively low sampling rates (typically 28-30 kHz) can otherwise introduce alignment artifacts.

### 2.3. Preprocessing for Feature Extraction

Following initial spike detection, robustly handling signal artifacts and overlapping spikes is essential for reliable single-neuron isolation, particularly in noisy clinical recordings. Our pipeline addresses this with a sequential, three-phase module designed to systematically clean data. This process consists of: (1) bundle-level artifact detection to identify widespread, non-neural artifacts, (2) within-channel artifact detection and collision detection to handle overlapping spikes on a single electrode, and (3) template-matching spike rescue to recover and decompose genuine neural signals.

Crucially, this three-phase module operates prior to feature extraction and dimensionality reduction, ensuring that both non-neural events and unresolved collisions do not contaminate the feature space used for clustering, where single unit templates need to be identified. This sequential arrangement substantially improves clustering outcomes through several mechanisms. It reduces the dimensionality of the clustering problem by eliminating irrelevant events, which improves computational efficiency. Furthermore, it enhances cluster separation by removing outliers and artifacts that can bridge otherwise distinct neural populations. This preprocessing also enables more accurate density estimation for clustering algorithms that rely on local density structure, as artifact-induced distortions in the feature space are minimized.

#### Bundle-Level Artifact Removal

The first phase targets widespread artifacts by analyzing spatial correlations across the entire electrode bundle (e.g., 8 channels). This method operates on a key principle: non-neural events arising from electrical interference or patient movement tend to create synchronous, high-amplitude signals across many channels at once. Conversely, genuine neural spikes are spatially localized, appearing in only one or a few nearby electrodes. This initial phase identifies and flags these spatially widespread events, distinguishing them from true neuronal activity.

The method operates by analyzing the signal across all channels in a bundle using a sliding time window of 0.5 ms. For each window, if spikes have been detected in 6 or more channels, the events are flagged as artifacts. Window length and number of channels can be adjusted to the specific recording setup and the characteristics of the artifacts encountered in human clinical recordings. By removing these artifacts early in the preprocessing pipeline, the method ensures that downstream steps, such as feature extraction and clustering, operate on cleaner data, ultimately enhancing the reliability of single-neuron isolation. The effect of this bundle-level artifact removal can be seen in Figure 4, which demonstrates the detection of non-neural artifacts within a bundle implanted in the left Amygdala of a subject, captured along with genuine neural spikes. In this case, on average, approximately 8.02% (min:2.84%, max: 23.14%, SD: 6.24%) of the acquired spikes across all the channels in the left Amygdala, recorded in 40 minutes, were bundle-level artifacts.

**Figure 4:**
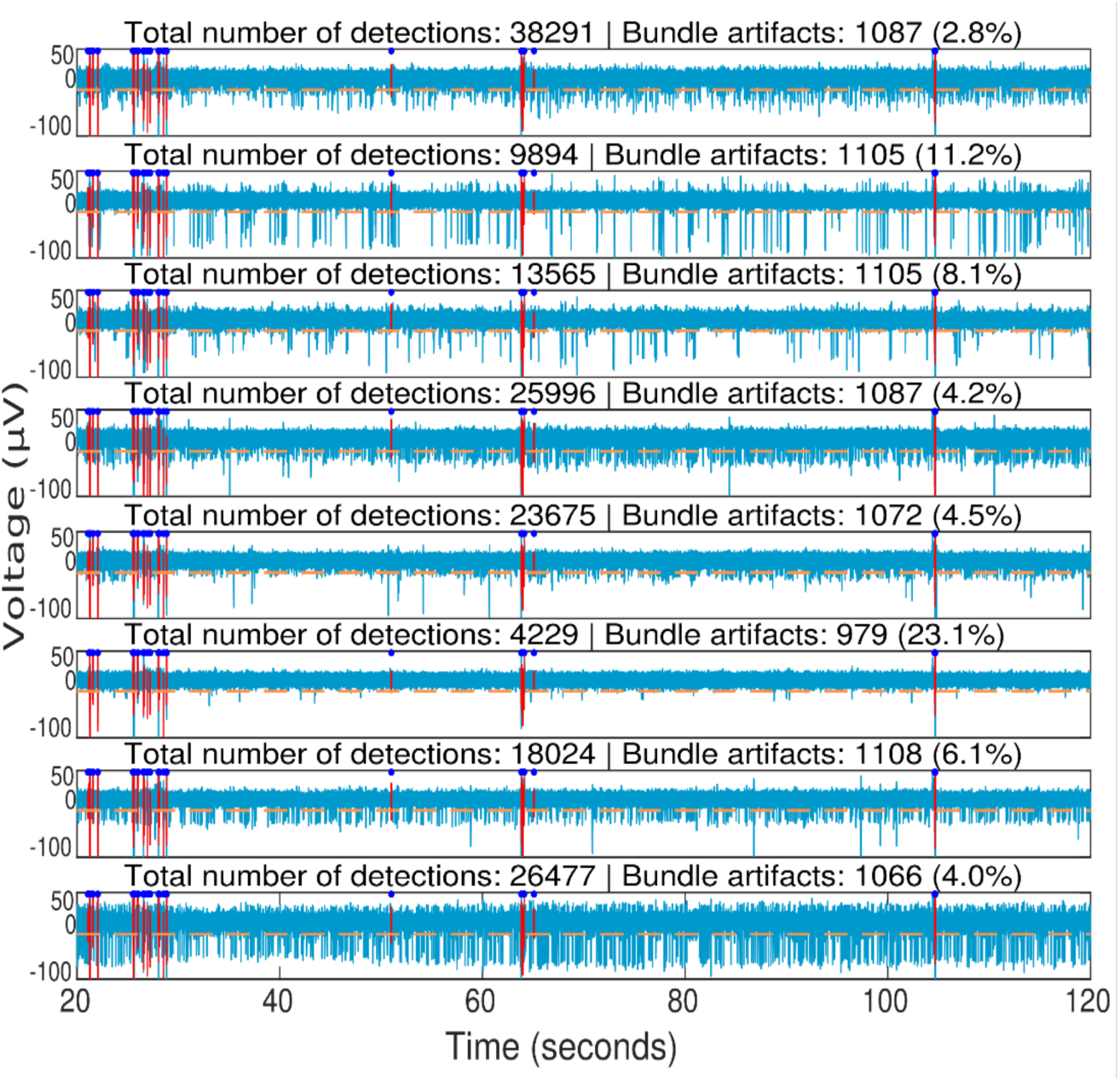
Bundle-Level Artifact Removal: The bandpass and notched-filtered signal (light blue) is processed by the Spike-Detection step, which detects spikes across 8 channels in the left Amygdala based on evaluated spike thresholds (orange dashed line). Time points of detected bundle-level artifacts are indicated above (dark blue dots), along with their corresponding waveforms (red). The pipeline accurately captures bundle-level artifacts while preserving genuine spikes, demonstrating minimal false positives. The figure also mentions the total number of detected spikes and the number of detected bundle-level artifacts for each channel. On average, approximately 8.02% of detected waveforms (min: 2.84%, max: 23.14%, SD: 6.24%) in the left amygdala were classified as bundle-level artifacts.

#### Within-Channel Artifact Rejection

The second phase of the pipeline operates on individual channels to identify and isolate potentially problematic waveforms. Rather than implementing immediate deletion, this phase utilizes a quarantine strategy, setting aside suspicious events for re-evaluation after the initial clustering is complete. This approach ensures that marginal signals are not prematurely lost, allowing for potential “rescue” once the broader context of neural activity is established.

Artifact detection is governed by a hierarchical decision logic, as illustrated in Figure 5, that evaluates waveforms based on amplitude, peak morphology, and temporal characteristics. Initially, the system filters out low-power noise by comparing the event amplitude to the global distribution; any signal falling below the 5th percentile is immediately quarantined. For signals exceeding this threshold, the system performs a peak-count analysis. Waveforms exhibiting a single, clear peak are passed directly to the final temporal validation. However, multi-peak waveforms, which often signify overlapping spikes or complex electrical interference, undergo a secondary prominence evaluation.

**Figure 5:**
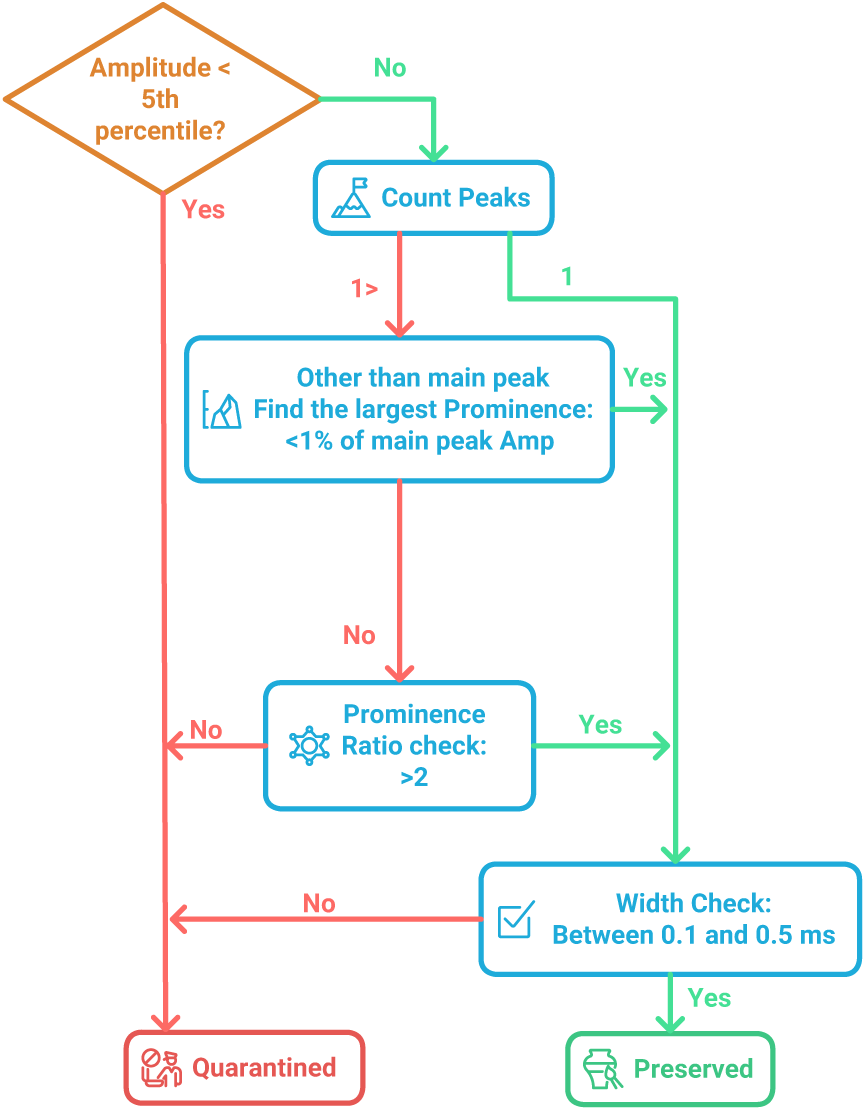
Hierarchical Decision Logic for Within-Channel Artifact Rejection. The flowchart depicts the sequential filtering process used to classify waveforms as either **Preserved** or **Quarantined**. The process begins with an amplitude thresholding (5th percentile), followed by a morphological assessment of peak counts. Multi-peak events are subjected to a two-stage prominence check: first, an absolute threshold (<1% of main peak amplitude) and second, a prominence ratio check (>2). Finally, waveforms must meet a temporal width constraint of **0.1 to 0.5 ms** to be accepted. Green paths indicate the criteria for signal preservation, while red paths denote the transition to the quarantine pool for events that fail to meet physiological or signal-integrity standards.

If the largest secondary peak has a prominence of less than 1% of the main peak’s amplitude, it is dismissed as negligible noise and the event proceeds. If the secondary peak is more substantial, the system calculates a prominence ratio. A ratio greater than 2 allows the event to continue through the pipeline, while events failing this check are flagged as artifacts. The final stage of the rejection logic is a physiological width check. Waveforms must possess a temporal duration between 0.1 ms and 0.5 ms to be preserved. Those falling outside this range, representing either high-frequency electrical shunts or slow-moving motion artifacts, are directed to the quarantine pool.

#### Template Matching and Waveform Rescue

The final phase of our collision detection pipeline employs a sophisticated two-pass template-matching procedure designed to recover quarantined waveforms and assign them to established clusters. For each established cluster, the algorithm constructs a template by computing the centroid (mean waveform) and establishing a spherical decision boundary around each cluster centroid with a radius 3𝜎*_τ_* where 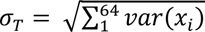 is the total variance estimate with 𝑣𝑎𝑟(𝑥*_i_*) denoting the variance at the *i*-th sample across the waveforms of a given cluster [16], [22]. The distance metric itself can be implemented using various approaches, including Euclidean distance, Manhattan distance, or other weighted distance measures, with the specific algorithm being less critical than the overall two-pass strategy.

The first pass focuses on single-cluster assignment, where each quarantined spike is systematically evaluated against all existing cluster centroids. The waveform is assigned to the cluster with the smallest distance to its centroid, provided this distance falls within the spherical decision boundary. This conservative approach ensures that only waveforms with high similarity to existing clusters are recovered, effectively rescuing single-unit spikes that were quarantined due to minor morphological variations, amplitude fluctuations, or threshold violations. Spikes that cannot be assigned to any single cluster during this pass proceed to the second phase for more sophisticated analysis.

The second pass implements a linear combination rescue strategy specifically designed to identify and recover overlapping spikes. This phase recognizes that when multiple neurons fire within a short temporal window, their action potentials can sum linearly, creating composite waveforms that do not match any single cluster but can be explained as combinations of multiple cluster centroids. The algorithm systematically evaluates all possible pairwise combinations of cluster centroids, testing whether each quarantined waveform can be explained as a linear combination of any two centroids at various temporal offsets. This process explores the possibility that the composite waveform represents two overlapping spikes from different neurons firing at slightly different times, where the linear combination of cluster centroids at various temporal offsets may explain the observed waveform.

When a waveform is successfully matched to a linear combination of two clusters, the algorithm performs spike decomposition to recover the individual neural events. The temporal offset that minimizes the distance between the composite waveform and the linear combination is identified as the optimal lag, representing the relative timing between the two overlapping spikes. Using this optimal lag, the algorithm performs waveform subtraction to isolate each spike’s contribution. The first cluster’s spike is recovered by subtracting the second cluster’s scaled centroid from the composite waveform, while the second cluster’s spike is recovered by subtracting the first cluster’s contribution. For each recovered waveform, the algorithm identifies the peak and determines the precise spike time, which may result in two spikes occurring within the physiological refractory period since different neurons can fire simultaneously without violating biological constraints.

The recovered spikes are then added to their respective clusters with their newly determined spike times and waveform data, effectively increasing the yield of genuine neural events from the original recording. However, the algorithm maintains strict quality control through conservative rejection criteria. If the closest linear combination among all possible cluster pairs still exceeds the distance threshold spherical decision boundary, the waveform is discarded as noise or an unresolvable artifact. This approach ensures that only high-confidence spike assignments are made, preventing the introduction of false positives that could compromise cluster integrity.

### 2.4. Experimental Task-Specific Processing

After removing collisions and artifacts, our pipeline incorporates an experimental task-specific processing module that is important for isolating neural activity directly relevant to the task in a real-world clinical environment. This module is designed to account for the myriad sources of variability and interference that are unique to human extracellular recordings in hospital settings, such as environmental disruptions arising from patient conversations, routine clinical care, mobile phone interventions, staff interruptions, meals, restroom breaks, or periods when the patient requires breaks within tasks. By leveraging detailed experimental metadata and stimulus timing information, the module ensures that only neural signals temporally and contextually aligned with the experimental protocol are retained for further analysis while rigorously excluding extraneous or confounding unrelated activity. To demonstrate this task-specific signal processing module, we present here the example of our Rapid Serial Visual Presentation (RSVP) task. Subjects sit in bed, facing a screen on which a sequence of 60 images is presented for 500 ms each picture, in pseudorandom order and with blanks to mark the beginning and end of a sequence. Each picture presentation is paired with a small square changing color for 50 ms, a transition that can be detected with a photodiode. The sequence of picture presentations is triggered with a keypress by the patient and lasts approximately 30 seconds [23]. Several of these sequences make a block, and several blocks comprise an experimental session, which lasts 30 to 40 minutes. In the context of this RSVP task, the pipeline’s experiment task-specific processing module starts with the extraction of the precise stimulus onset times from the photodiode channel. Thereafter, it matches the detected onsets to the expected number of stimuli based on the experimental design, corrects for any discrepancies, and generates a visual and quantitative check on how accurately the photodiode-based event extraction has identified stimulus onsets. The module is also critical for segmenting neural data according to precise experimental events, such as the onset and offset of each sequence.

This photodiode-based temporal segmentation approach, illustrated in Figure 6, enables our pipeline to focus exclusively on neural recordings during active experimental periods, effectively isolating task-relevant neural activity from potential artifacts. The top panel shows the full recording session over approximately 58 minutes, where the clear gaps between stimulus blocks (visible as intervals with no red photodiode signal) were associated with extended breaks requested by the subject during the task, completing only 3 task blocks over an hour, as opposed to the typical 5 blocks within approximately 30 to 40 minutes. During these breaks, there were significant movement artifacts that were recorded. The middle panel demonstrates how these precisely timed experimental epochs, marked by the sequence onset and offset (black dashed lines), ensure that spike sorting algorithms operate only on neural data that is temporally aligned with the cognitive task, thereby eliminating potential contamination from irrelevant neural activity and maximizing the likelihood of detecting stimulus-locked responses. The bottom panel of Figure 5 illustrates the sub-millisecond precision of photodiode-detected stimulus onsets, with sharp onset transitions providing the temporal accuracy necessary for reliable neural signal processing.

**Figure 6:**
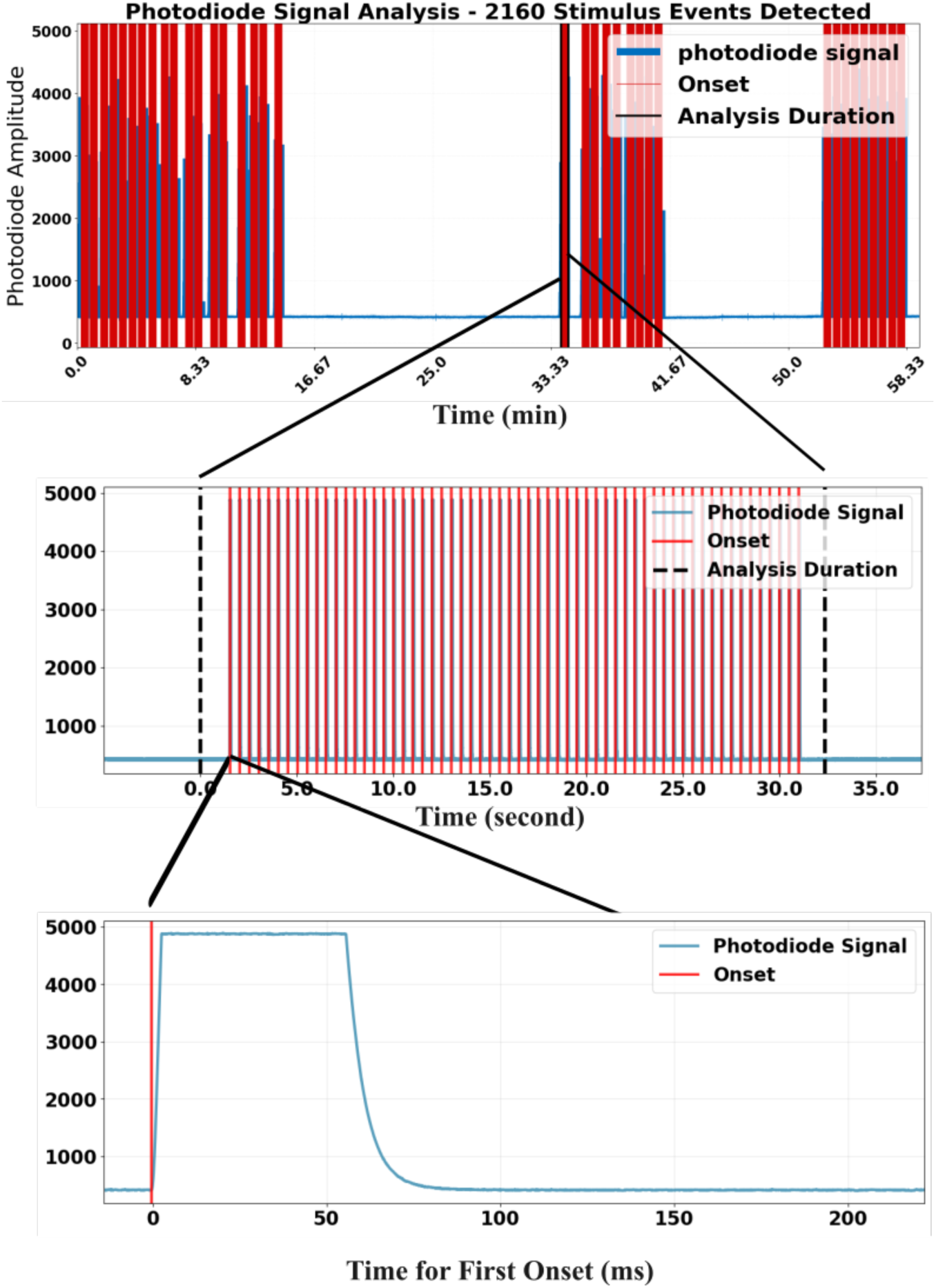
Photodiode-based data segmentation: The figure demonstrates photodiode-based stimulus event detection and temporal segmentation for the RSVP experimental paradigm, with 2160 stimulus events detected across the complete recording session. The top panel shows the photodiode amplitude over the full experimental timeline spanning approximately 58 minutes, revealing three blocks with two long gaps in between, each with distinct epochs (sequences) where stimulus presentation occurs (red photodiode signal peaks)The gaps between sequences can be precisely identified, enabling the pipeline to focus exclusively on neural recordings during active experimental periods while excluding contaminated data from non-task intervals such as patient rest periods, clinical interruptions, or within-session breaks. The middle panel provides a focused view of the first sequence in the second task block, illustrating the 2Hz stimulus presentation pattern characteristic of this RSVP paradigm, with sequence onset and offset (black dashed lines) clearly demarcating the temporal window for neural data extraction. The bottom panel shows a detailed view of the first stimulus onset detection. The 50 ms long square changing color in sync with picture onset generates transient changes in the photodiode signal that can be accurately detected, providing sub-millisecond timing precision. This temporal precision, combined with the clear segmentation of task-relevant periods, ensures that subsequent spike sorting and neural analyses are performed exclusively on data collected during well-defined experimental epochs, thereby eliminating potential confounds from extraneous neural activity unrelated to the cognitive task and maximizing the SNR for stimulus-locked neural responses.

Events not associated with picture presentation (including sequence onset and offset) are detected by a Data Acquisition (DAQ) system, which can be used as a backup timing source to the photodiode. These DAQ events are time-stamped in the same data stream as the neural recordings, providing a direct way to align neural activity with experimental events. While the DAQ serves as a useful backup, the photodiode is the preferred method for extracting picture presentation times, as it is directly synchronized with actual frame changes on the monitor. In contrast, relying on the DAQ alone can introduce timing inaccuracies because a standard monitor with a 60 Hz refresh rate may result in a delay of approximately 17 milliseconds for each missed frame, potentially affecting the latency estimation of neural responses. Hence, the pipeline ensures that only neural activity during well-defined experimental epochs is retained for analysis, while contaminated data from irrelevant periods is filtered out.

The comparative analysis presented in Figure 7 demonstrates the critical importance of task-specific clustering for isolating genuine neuronal units from the complex signal environment of clinical recordings. As shown in the figure, while both approaches detect multiple clusters, only a single cluster from each method represents an actual neuronal unit, Cluster 2 from task-specific processing and Cluster 6 from standard processing, with all other detected clusters representing noise, artifacts, or contaminated signals that must be appropriately excluded from neuronal analyses. The waveform density plots in Figure 7 provide compelling visual evidence of the superior unit quality achieved through task-specific clustering, with Cluster 2 displaying a remarkably narrow and well-defined density distribution compared to the broader, more diffuse pattern of Cluster 6 from standard processing. This visual difference is quantitatively supported by the dramatic improvement in key quality metrics: the task-specific approach yields a neuronal unit with zero refractory period violations (ISI<3.0ms: 0.0% vs 0.67%), substantially higher SNR (SNR: 4.69 vs 3.26), and higher firing rate (0.72Hz vs 0.63Hz). Critically, the task-specific unit represents 36% of all detected events compared to only 7% for the standard processing unit. This indicates that task-specific clustering creates a cleaner signal where genuine neuronal events form a dense, coherent cluster, whereas standard processing generates many noisy waveforms that fragment the feature space and make it difficult to isolate the true neuronal signal from spurious detections. The presence of refractory violations and lower SNR in the standard approach confirms contamination from noise or multi-unit activity. Figure 7 thus illustrates how task-specific clustering effectively separates genuine neuronal signals from the complex mixture of biological and technical noise inherent in clinical intracranial recordings, providing a more reliable foundation for subsequent neuroscientific analyses by prioritizing signal quality over quantity.

**Figure 7:**
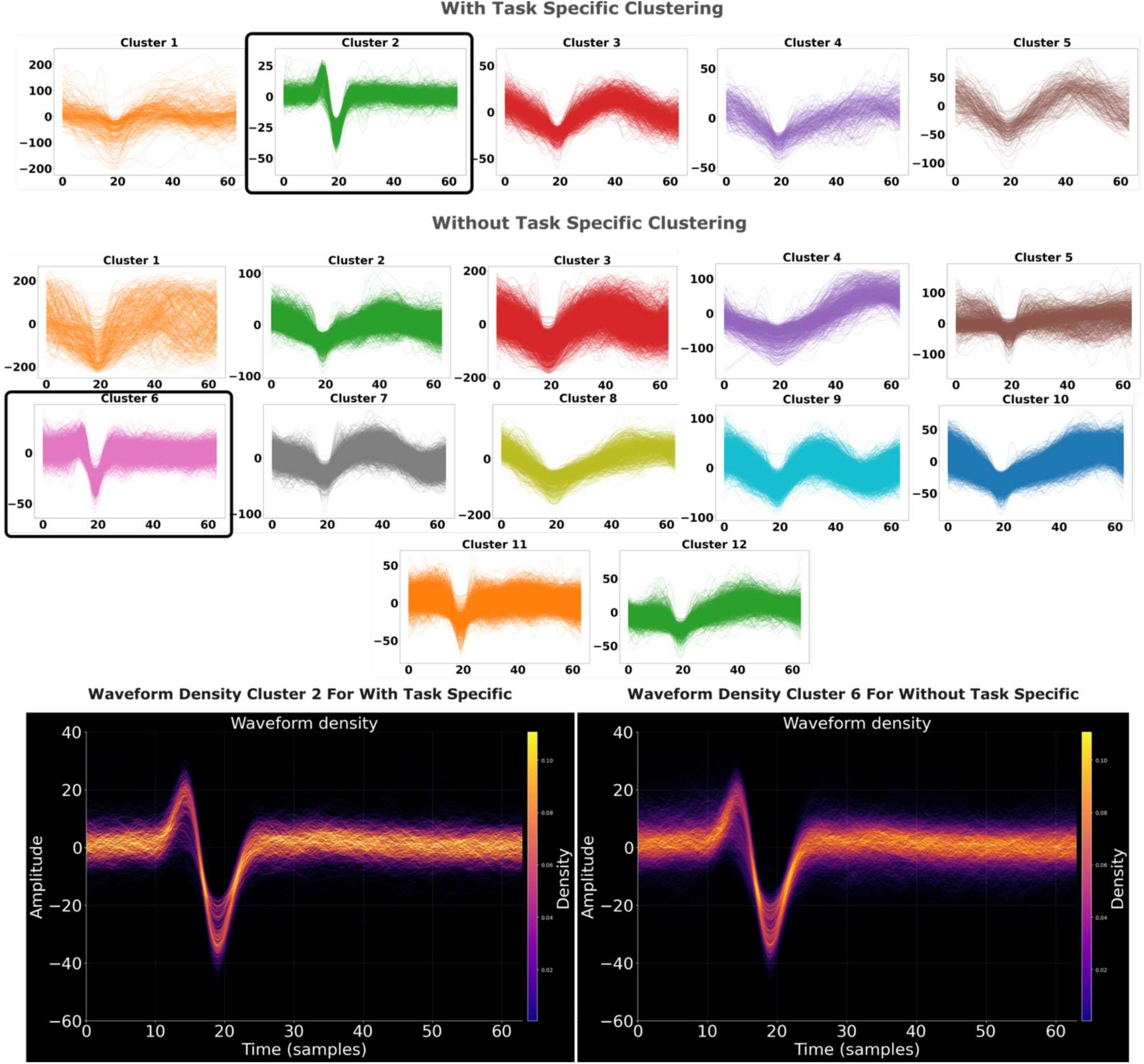
Separate Task. The top row shows results from task-specific clustering, which identified 5 clusters, with only Cluster 2 representing a genuine neuronal unit with well-defined waveform morphology. The remaining clusters appropriately isolate noise and artifacts. In contrast, standard clustering (middle and bottom rows) produced 12 clusters with only Cluster 6 representing the actual neuronal unit, while 12 clusters contained mixed noise, artifacts, and contaminated signals. The waveform density plots (bottom) highlight the key difference: Cluster 2 from task-specific processing (left) shows a narrow, well-defined density distribution with sharp temporal features, while Cluster 6 from standard processing (right) exhibits a broader, more diffuse pattern indicating greater waveform variability. Quantitative metrics confirm the superior quality of task-specific clustering: Cluster 2 demonstrates zero refractory period violations (ISI<3.0ms: 0.0% vs 0.67%), higher SNR (SNR: 4.69 vs 3.26, slightly higher firing rate (FR: 0.72Hz vs 0.63Hz), and represents a much larger proportion of detected events (36% vs 7% of total events). This demonstrates that task-specific clustering more effectively isolates genuine neuronal signals from complex biological and technical noise. In addition, waveform density plots (bottom panels) reveal that task-specific Cluster 2 exhibits higher peak density (max density: 0.111) compared to standard processing Cluster 6 (max density: 0.083), indicating more consistent waveform shapes and tighter clustering of genuine neuronal events.

### 2.5 Feature Extraction and Dimensionality Reduction

Following spike detection and artifact removal, extracted waveforms reside in a high-dimensional space where each dimension corresponds to a sampling point. Detected spike waveforms typically comprise 40-65 sample points (at ∼30 kHz sampling rate), creating high-dimensional representations where the complexity of clustering calls for dimensionality reduction.

The main issue is how to choose the minimal set of features that yields the best discrimination between neuronal units [9]. High-dimensional data processing requires substantial computational resources, which becomes particularly problematic for real-time or large-scale human recordings in clinical settings with limited computing infrastructure. As dimensionality increases, data becomes increasingly sparse in the feature space, a phenomenon known as the curse of dimensionality, which compromises distance-based clustering algorithm performance and reduces statistical power. The goal is to keep only the features that help classification, given that eliminating inputs dominated by noise can improve clustering outcomes [24].

Our pipeline addresses these challenges by transforming raw waveforms into lower-dimensional representations that emphasize discriminative characteristics while suppressing noise-related variability. The framework allows users to select from multiple dimensionality reduction techniques based on their specific recording characteristics, experimental requirements, and computational constraints. This adaptability is essential for human recordings, where signal quality, neural population characteristics, and noise profiles vary substantially across subjects and recording sites.

Principal Component Analysis (PCA) serves as a classical dimensionality reduction approach, identifying orthogonal directions of maximum variance in the data [25], [26]. For spike sorting applications, we compute the principal components from aligned spike waveforms and extract the first 2-3 components that typically account for most of the total variance. This approach provides a computationally efficient representation that preserves the global structure of the data while significantly reducing dimensionality. While computationally efficient, PCA has limitations for spike sorting. It assumes linear relationships between features and cannot capture the complex non-linear patterns that often distinguish different neuron types. Additionally, PCA optimizes for maximum variance preservation rather than maximum class separability, meaning that directions of high variance may not correspond to the most discriminative features for distinguishing between neuronal units.

Wavelet-based feature extraction simultaneously captures both temporal and spectral characteristics of spike waveforms, offering advantages for human microwire recordings [22], [27]. Wavelets are effective at representing localized waveform features that differentiate neurons, with the wavelet transform decomposing waveforms at multiple scales to detect both broad and fine waveform differences. The implementation includes automatic selection of discriminative wavelet coefficients based on the Kolmogorov-Smirnov (KS) test for non-normality [16], and another one using a novel **Gaussian mixture model (GMM)** approach to identify multimodal distributions of wavelet coefficients that likely represent multiple distinct neuronal clusters [28]. Good separability can then be achieved by selecting multiple wavelet coefficients providing non-redundant information. To this end, we will also remove redundant coefficients that would help to separate the same clusters.

#### GMM Feature Extraction

To utilize sparse or noisy data, a data-driven feature extraction is needed to judiciously select coefficients and checks for redundant information. Previous works involving GMM use for feature extraction used similar metrics from we will describe for their feature extraction but did not combine the metrics or select them in a data-driven way[28]. We expand on this methodology to combine two calculated metrics, then selecting coefficients, and removing redundancies. The KS test solely looks for non-unimodality, where this GMM function gives us more insight into the makeup of the waveforms, emphasizing multimodal coefficient waveforms over skewed waveforms, heavy-tailed waveforms, or other general unimodal but non-normal waveforms.

Following wavelet decomposition, each coefficient, spanning all detected spikes, is modeled using a GMM comprised of 16 sub-Gaussians. We first identify the **“main-mixture”** by locating the peak of the probability density function (PDF) and including all sub-Gaussians that fall within the half-prominence of that peak. A sub-Gaussian is statistically defined as part of this main mixture if its mean ± 2 standard deviations remain within the half-prominence boundary. This main mixture is characterized by its mean 𝜇_&_ and standard deviation 𝜎_&_, as visually represented by the blue waveforms and shaded region in **Figure 8A**.

**Figure 8:**
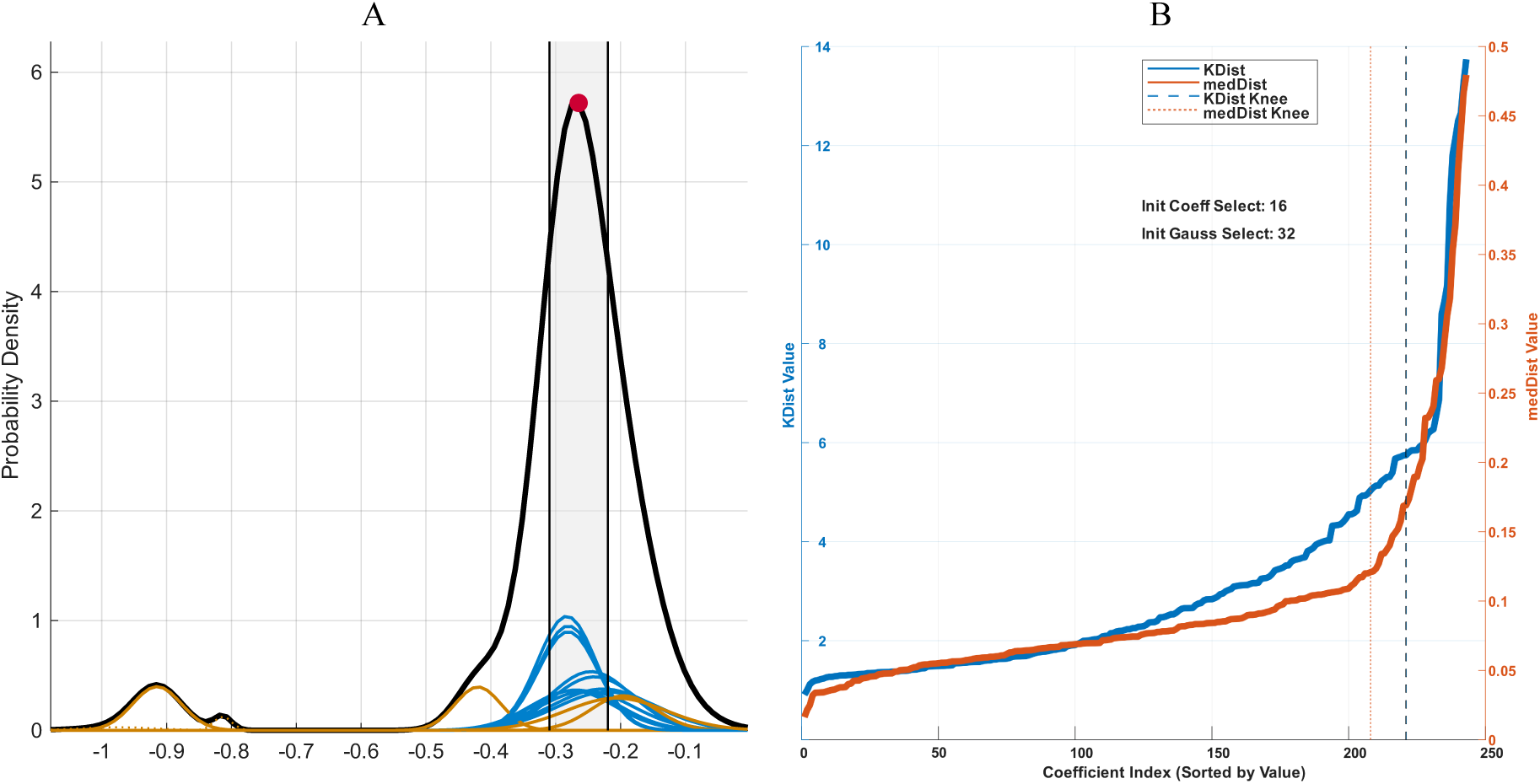
GMM-Based Coefficient Selection Metrics. **(A) PDF and GMM Decomposition.** The black curve represents the aggregate PDF of a single wavelet coefficient. The underlying sub-Gaussians are color-coded: blue curves represent the “main-mixture” sub-distributions located within the shaded half-prominence region of the global peak (red dot), while orange curves denote secondary or outlier distributions. **(B) Comparison of Selection Metrics.** 𝐾_dist_ (blue line, left axis) and 𝑀𝑒𝑑𝐷𝑖𝑠𝑡_i_ (orange line, right axis) values are plotted across all coefficient indices, sorted by value. Vertical dashed and dotted lines indicate the “knee” points for 𝐾_dist_ and MedDist_i_ respectively, which serve as data-driven thresholds for initial coefficient and Gaussian selection (indicated here as 16 coefficients and 32 Gaussians). The complementary nature of these curves allows for a more robust selection of multimodal features compared to single-metric approaches.

To evaluate the distribution of these coefficients, we calculate two complementary metrics:

1. 𝐾*_dist_*: This metric quantifies the isolation of an individual Gaussian from the main mixture by calculating the normalized distance between their means:

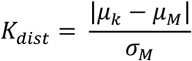

While effectively identifies Gaussians isolated from the primary signal, it is less sensitive to narrower distributions.

2. MedDist*_i_* : To address the limitations of 𝐾*_dist_*, we introduce a second criterion that utilizes a pairwise calculation to assess the distance between sub-Gaussians, accounting for their respective weights 𝛼. It is defined as:

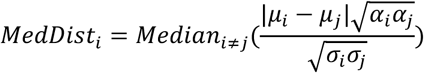

This metric is particularly robust for identifying narrow distributions but may overlook groupings of Gaussians situated outside the primary mixture.

The relationship between these metrics across the coefficient indices reveals a clear “knee” or elbow point for both curves. By identifying these transition points (indicated by dashed lines in **Figure 8B**), we can objectively select the most informative coefficients while discarding those that contribute primarily to background noise.

To integrate these metrics into a robust selection criterion, we utilize a bivariate decision framework that maps both values into a shared feature space. This process begins by identifying the “knee” points for both metrics in their respective distributions, as illustrated by the dotted lines in **Figure 8B**. These threshold values are then used to establish a reference point in a two-dimensional space where *𝐾_dist_* is plotted against *MedDist_i_*. The intersection of these metric knees, represented by the yellow circle in **Figure 9A**, serves as the primary anchor for our decision boundary.

**Figure 9:**
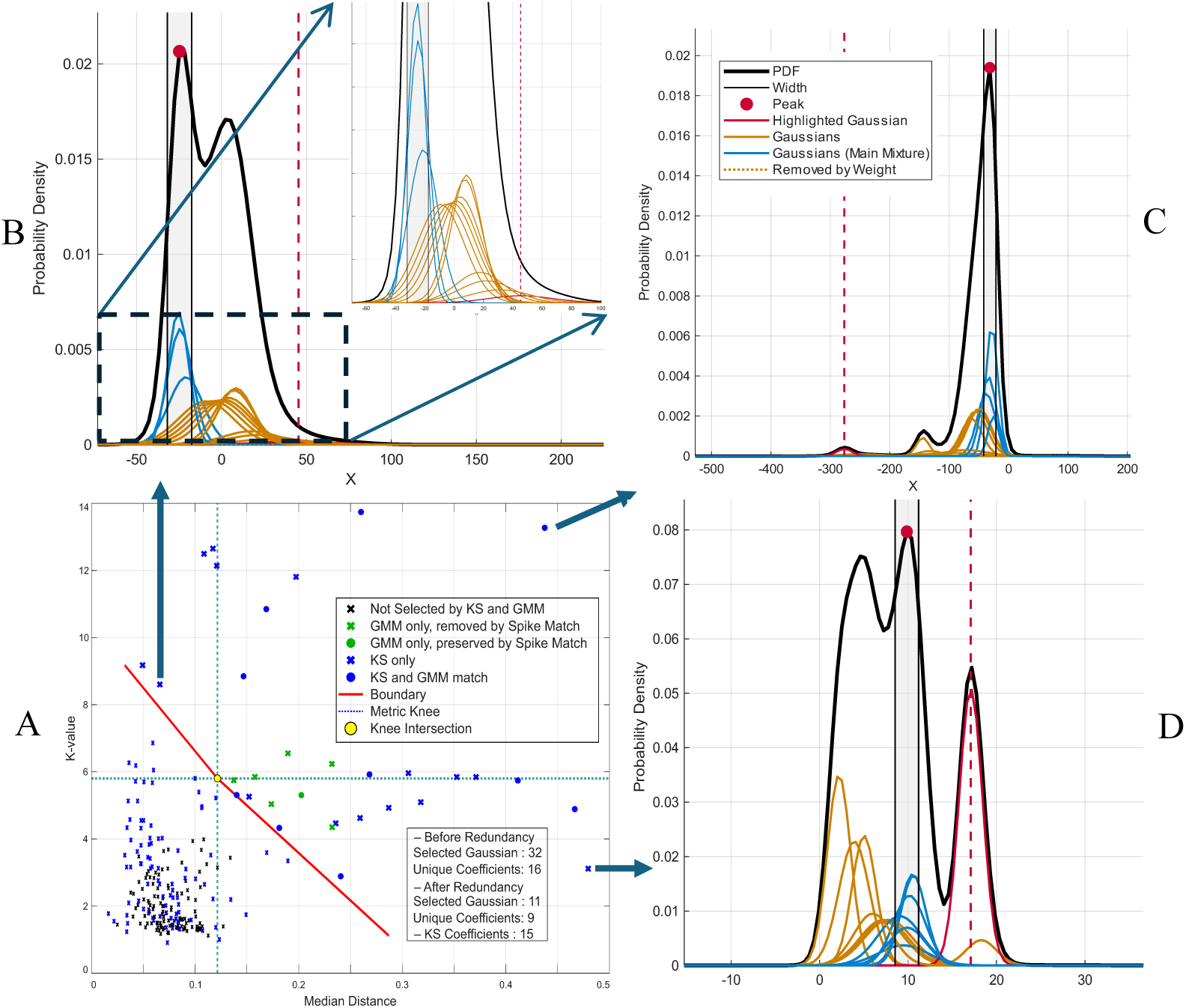
Boundary Line Creation. **A.** The 2D plot with created boundary lines are seen in these figures. Values above the boundary line are considered passing and values below are excluded. This plot also compares the KS method to the GMM, showing values selected by KS only, KS and GMM, GMM only, and GMM only but was redundant. In seeing these we can see how we remove a lot of redundant information that the KS method selected, as well as bring unique information that the KS method missed, while still having the GMM method check itself for redundancies. In figures **B-D** we see how different locations in the 2D boundary space manifest very different looking waveforms, emphasizing the need for both Kdist and medDist metrics to be included in feature extraction, the main mixtures are identified by the shaded regions and their sub-gaussians are blue, the sub-gaussians outside are orange, and the selected gaussian of focus is red. **B** this sub figure shows a zoomed in view of the gaussians, allowing us to see why a Kdist value would be large, but medDist would be low, and that a gaussian with a relatively decent weight could be weighted highly due to its width.

A linear boundary is constructed from this knee intersection point toward the distribution’s extremities: the point with the highest *𝐾_dist_* at the lowest *MedDist_i_*, and the point with the highest *MedDist_i_* at the lowest *𝐾_dist_*. To prevent the boundary from being skewed by statistical anomalies, extremity points, defined as those residing in the top 10% distance-wise from the knee intersection, are excluded during the line-fitting process. Waveforms and coefficients located above this red boundary line are preserved, while those below are discarded as noise or redundant information.

This multi-metric approach offers a more nuanced selection than traditional methods like the Kolmogorov-Smirnov (KS) test. As shown in **Figure 9A**, the GMM-based boundary identifies physiologically relevant information that the KS method misses, while simultaneously filtering out a significant amount of redundant data that the KS method erroneously includes. The necessity of using both metrics is further validated in the morphological profiles shown in **Figures 9B–D**. For example, **Figure 9B** illustrates a waveform where a specific Gaussian exhibits a high 𝐾*_dist_* but low *MedDist_i_*. Despite its isolation from the main mixture, its specific weight and width indicate it is a significant feature rather than noise, a distinction enabled by the dual-metric evaluation.

Following the boundary line creation all points above the boundary line are selected and move on for further processing. The goal of the next and final processing step is to remove redundant information from the gaussian-coefficient pairs selected by the boundary line. Obviously, a coefficient with multiple gaussians above the line will be easy enough to find, however, making sure that a set of spikes represented by each gaussian are unique in the selected gaussian-coefficient pairs is a novel and unique aspect of the GMM model that we will use to avoid redundant coefficient selection. To do this first we find the set of the spikes, *s_k_*, represented by a given gaussian. This is defined by the wavelet range [𝜇_*_ ± 2𝜎_*_], where µ_k_ and σ_k_, are defined by a sub-gaussians mean and standard deviation, and the spikes that have a value in the given range for the coefficient paired with the gaussian are included in the set. Then the overlap coefficient is determined for all selected gaussian-coefficient in pairwise fashion, or *Overlap* (k, ℓ) = 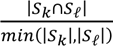, Finally, redundancies are removed using a Pearson correlation on each row of the overlap matrix and removing gaussian-coefficient pairs with greater than 0.9 correlation. The resulting coefficients are the features used for clustering. When compared to the KS extraction method, we result in fewer coefficients for the same number of neurons but still have a trend increasing the coefficients as the number of neurons increases as seen in figure 10. This implies that our model reduces the redundant coefficients and makes more room for unique but less represented data to appear, however further validation is needed to confirm this.

**Figure 10:**
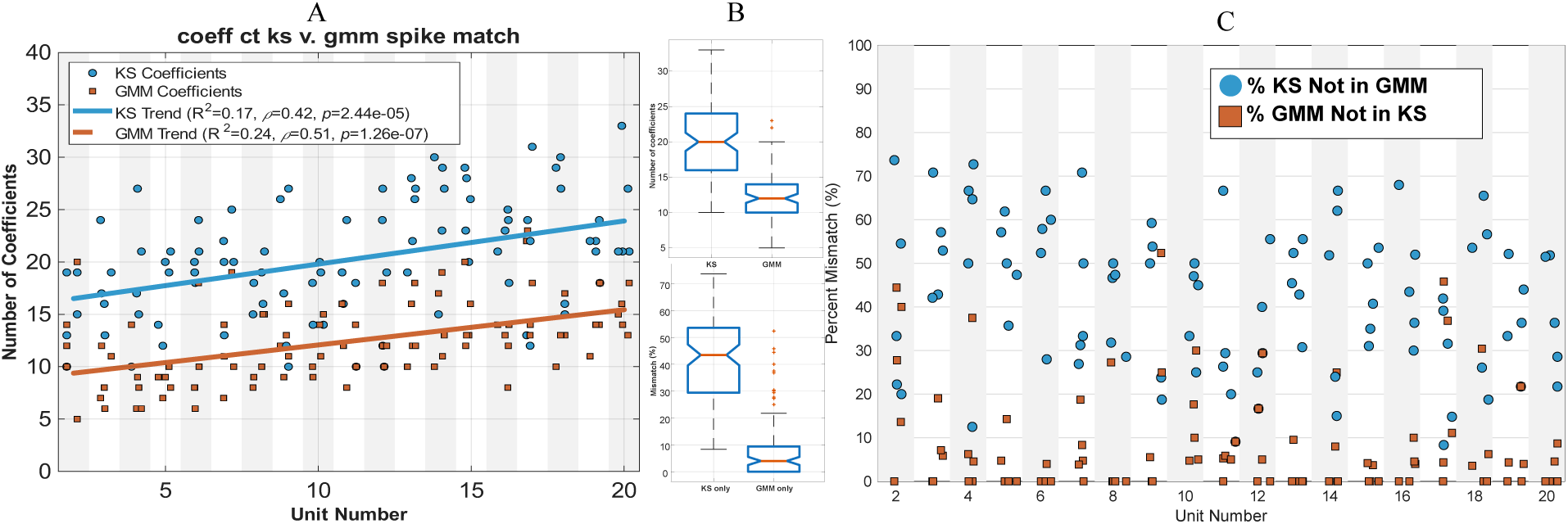
Comparison of GMM to KS. **A.** This figure shows that both GMM and KS have a positive trend with the number of simulated neurons (5 simulations per unit), but that the GMM model tends to have less coefficient overall, implying a more selective feature extraction process. **B-C.** These two figures represent the overlap of coefficients between the GMM and KS methods. Mismatch is the percentage of a given coefficient not included in the other, match is the percentage of coefficients overlapping between GMM and KS. Note that GMM had less variation in match and mismatch (**B**) and tended to have lower amounts of coefficients not in the other, further implying the redundancy removal worked. Percentages are calculated over the whole number of coefficients between both methods.

### 2.6. Clustering Algorithms

The critical task of assigning individual spikes to putative neuronal sources is accomplished through clustering algorithms. Clustering represents one of the most challenging aspects of the spike sorting pipeline, particularly in human microwire recordings, where variable noise profiles, non-stationary signals, and complex neuronal dynamics introduce significant complexity. Our pipeline implements multiple complementary clustering approaches, allowing users to select the most appropriate algorithm for their specific recording conditions and experimental requirements.

#### Available Methods

Our pipeline also incorporates superparamagnetic clustering (SPC), a physics-inspired clustering algorithm that models the clustering problem as a spin glass system [22]. The algorithm operates across different “temperature” parameters to identify natural cluster boundaries, with the super-paramagnetic regime providing optimal separation of neuronal units. SPC is particularly advantageous for human microwire recordings due to its ability to handle irregular cluster shapes and varying densities without distributional assumptions, capabilities that are especially important given the heterogeneous nature of neural signals in clinical recordings. To integrate SPC into Python-based pipelines, we created a Python wrapper that calls the C code directly in-memory, making it faster than the standalone executable. The wrapper is easy to install, fully cross-platform, and integrates seamlessly into existing workflows[29].

We also implement ISOSPLIT, a clustering algorithm specifically designed for neural spike sorting that operates on cluster isolation principles rather than density estimation [30]. The algorithm iteratively tests partition boundaries between cluster pairs using one-dimensional projection tests, automatically determining cluster numbers and handling complex geometries typical of neurophysiological data. This approach is particularly well-suited for human recordings where the number of distinct neurons is unknown a priori and signal quality can fluctuate within sessions.

Our pipeline also incorporates DBSCAN [31], a density-based clustering approach that defines clusters as continuous high-density regions separated by lower-density areas. DBSCAN intrinsically handles noise as outliers while identifying clusters of arbitrary shapes, making it particularly well-suited for neural data where cluster boundaries may be irregular and noise contamination is common. The algorithm requires two key parameters: the neighborhood radius (epsilon) and the minimum number of points required to form a dense region, allowing it to automatically determine the number of clusters without prior specification.

### 2.6. Quality Assessment

Reliable interpretation of neural data crucially depends on the accurate identification of single-unit activity. In human microwire recordings, where neural signal quality can vary substantially due to clinical constraints and recording longevity, rigorous quality assessment becomes particularly important. Our pipeline incorporates comprehensive quality assessment tools that enable objective evaluation of sorting results and facilitate meaningful comparisons across subjects, brain regions, and recording sessions.

#### Quality Metrics Implementation

Our pipeline leverages and extends the quality metrics framework from SpikeInterface [19], [20] to provide a robust assessment of sorted units. This implementation includes a comprehensive suite of established quality metrics that collectively characterize different aspects of sorting performance. This evaluation begins by characterizing basic unit features such as SNR on unit templates, average firing rate, and the fraction of time the unit is present (presence ratio). To assess stability and potential spike loss, we compute the amplitude cutoff from the unit’s amplitude histogram and track waveform shifts over time using maximum and cumulative drift metrics. Spike train quality is critically evaluated by calculating the rate of Inter-Spike Interval (ISI) violations to identify biologically implausible firing patterns. Lastly, cluster quality and isolation are quantified using a battery of established scores, including isolation distance, L-ratio, D-Prime, nearest-neighbor contamination, and the silhouette score, allowing for a thorough, multi-faceted validation of the sorting output.

#### Sorting Agreement and Benchmarking

Given the inherent challenges in establishing ground truth for human microwire recordings, our pipeline implements comprehensive sorting agreement and benchmarking tools that enable systematic comparison of different spike sorting approaches. The core of this implementation utilizes pairwise sorter comparison methods that apply the Hungarian algorithm [32] to find optimal matching between units detected by different sorters, using a 0.5 (ms) coincidence window to account for slight temporal alignment differences [19]. This process generates a unit match map and calculates an agreement score ranging from 0 to 1 for each putative neuron, providing an objective measure of sorting consistency across algorithms.

For studies employing multiple sorting algorithms, our pipeline extends this approach to multi-sorter consensus analysis, which constructs a graph representation of unit relationships across sorters and extracts consensus subgraphs for cases where three or more sorters agree. This approach is particularly valuable for human recordings where the absence of ground truth makes algorithm selection challenging, as it leverages the complementary strengths of different sorting approaches while mitigating their individual weaknesses. When ground truth data is available, typically through simulation or benchmark datasets with known spike times, the pipeline supports direct comparison between sorting results and ground truth, classifying spikes as true positives, false positives, or false negatives, and computing standard performance metrics including accuracy, precision, recall, miss rate, and false discovery rate.

Based on these comparison metrics, our pipeline categorizes detected units using standardized criteria: well-detected units (agreement ≥ 0.8), false-positive units (agreement < 0.2), redundant units (multiple detections of the same neuron), and over-merged units (inappropriate combining of spikes from distinct neurons). This categorization provides an objective framework for evaluating sorting quality and facilitates comparison across studies and laboratories. For comprehensive evaluation across multiple datasets and algorithms, our implementation includes a benchmark platform that can batch-run multiple sorters and aggregate results into structured data tables suitable for generating performance dashboards and statistical analyses.

#### Response-Profile Validation

Beyond waveform and spike train statistics, our pipeline incorporates response-profile validation methods specifically designed for human cognitive neuroscience applications. This approach leverages the expectation that many neurons in regions such as the medial temporal lobe exhibit selective responses to specific stimuli or cognitive conditions, providing an additional, functionally relevant dimension for quality assessment.

For validating spike sorting in datasets with specific experimental manipulations, the pipeline supports blinded analysis workflows where sorting is performed without knowledge of experimental conditions, followed by statistical validation of expected response patterns in the sorted units. This approach is particularly valuable for confirming the biological plausibility of detected units through their functional characteristics rather than waveform features alone.

The integration of these functional validation methods with traditional quality metrics provides a multi-dimensional quality assessment framework, particularly suited to human cognitive neuroscience recordings, where the scientific value of detected units ultimately depends on their ability to reveal meaningful neural correlates of cognitive processes and behaviors. This comprehensive approach maximizes confidence in the biological validity of sorted units while providing transparent quality metrics that can be reported in publications to facilitate interpretation and replication.

This response-profile validation approach, illustrated in Figure 11, provides crucial functional confirmation that our sorted units represent genuine neurons with distinct stimulus selectivity rather than contaminated signals or artifacts. As demonstrated in panel A of Figure 11, Cluster 4 (red-boxed, corresponding to the red waveform trace in panel B) exhibits clear stimulus-locked responses with robust firing rate increases to pictures of cats, while remaining completely silent during presentation of Trump pictures. Conversely, Cluster 3 (corresponding to the blue waveform trace) shows selective responses exclusively to pictures of Trump within the same experimental session, with no detectable firing during cat picture presentations. This mutual exclusivity in stimulus responses, where each cluster responds robustly to one stimulus category while remaining silent to the other, provides compelling evidence that our clustering algorithm successfully isolated two distinct neurons with complementary selectivity profiles.

**Figure 11:**
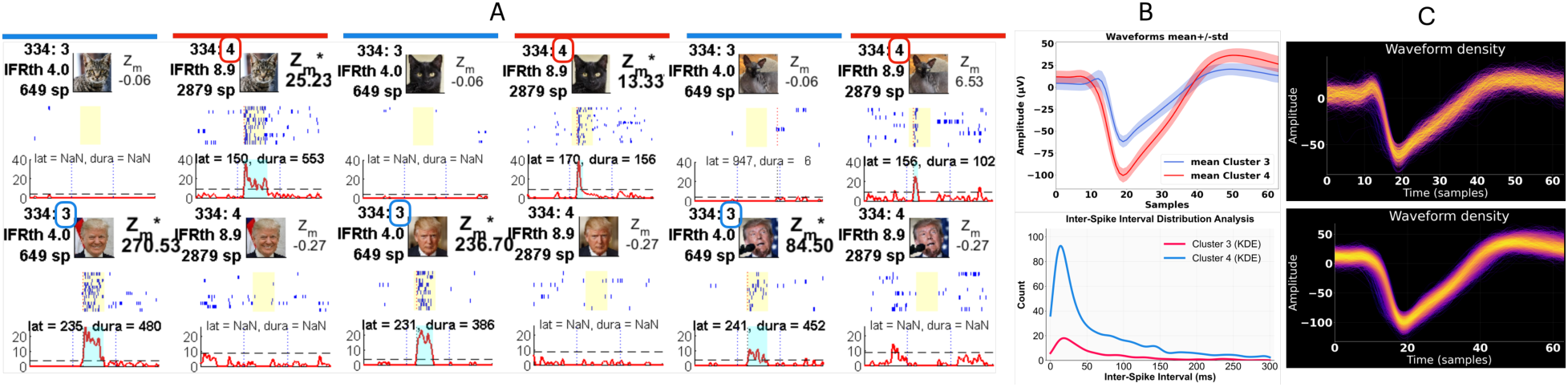
Response-profile validation confirms biological validity of sorted units through stimulus-selective neural responses. Panel A presents the critical functional validation, showing stimulus-evoked firing rate responses through peristimulus time histograms (PSTHs) that demonstrate clear functional dissociation between sorted units. The red-boxed examples highlight Cluster 4, which exhibits robust stimulus-locked responses to one category of pictures while remaining completely silent during presentation of different stimuli. Conversely, Cluster 3 shows selective responses exclusively to the alternative stimulus category with no detectable firing during the first stimulus type. Panel B(top) displays the mean waveforms ± standard deviation for Clusters 3 and 4, confirming distinct waveform morphologies corresponding to the functionally validated units. Panel B(bottom) shows the inter-spike interval (ISI) distributions with kernel density estimation (KDE) for both clusters, demonstrating appropriate refractory period characteristics that confirm biological validity. Panel C(top) presents the waveform density plot for Cluster 3, while Panel C(bottom) shows the waveform density plot for Cluster 4, both revealing the temporal stability and amplitude consistency of spike waveforms across the recording session. This multi-dimensional validation approach demonstrates that sorted units not only exhibit the expected functional properties of neurons in cognitive brain regions through distinct stimulus selectivity but also meet traditional spike sorting criteria through high-quality waveform characteristics and proper ISI distributions. The consistency of stimulus-specific functional responses across repeated presentations, combined with stable waveform properties and biological ISI patterns, ensures that detected neural signals represent genuine single-unit activity rather than noise or multi-unit contamination, thereby validating their suitability for neuroscientific analyses of stimulus-response relationships.

The consistency of these stimulus-specific responses across repeated presentations, combined with the high-quality waveform characteristics shown in panels B and C, validates that these sorted units represent biologically meaningful single neurons rather than noise or multi-unit contamination. This functional validation is particularly important in human intracranial recordings, where the clinical environment introduces numerous potential sources of signal contamination.

Beyond demonstrating the effectiveness of our clustering approach, the functional diversity observed between nearby neurons highlights the remarkable heterogeneity of neural populations in the human cortex. Figure 12 illustrates this diversity even more dramatically, showing two neurons isolated from the same microwire that exhibit vastly different selectivity profiles despite being within 100 microns of each other in the human fusiform cortex. While one neuron responds broadly to 79.9% of presented stimuli with extremely low selectivity, the neighboring neuron shows highly selective responses to only 2.2% of the same stimulus set. This striking difference in response patterns - from broadly responsive to highly selective - demonstrates that our sorting algorithm successfully captures the full spectrum of neural selectivity present in human recordings. The non-categorical nature of the selective responses further underscores the complex functional organization of the human cortex, where neurons with dramatically different coding properties can coexist in close spatial proximity. These findings validate not only our technical approach but also reveal the rich functional diversity that makes human single-unit recordings uniquely valuable for understanding cortical computation.

**Figure 12:**
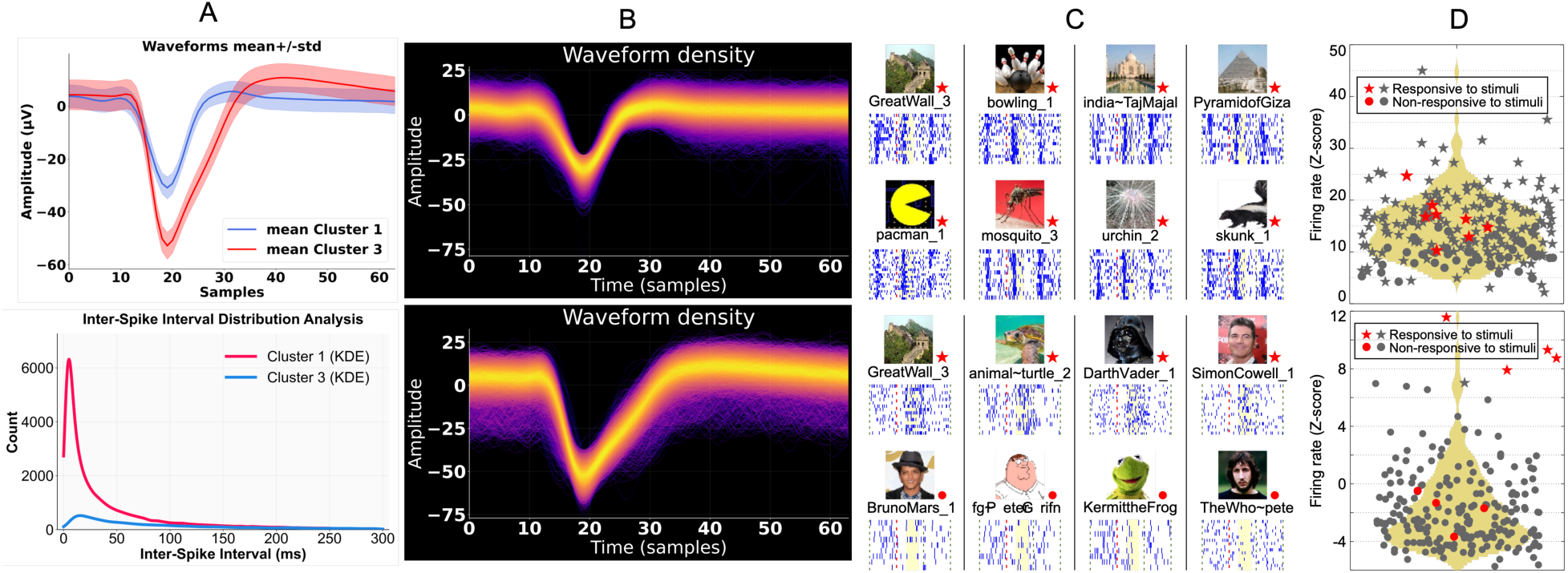
Very different selectivity can be seen across neurons within 100 microns. Two different neurons were isolated from a single wire. **A(top)**. Mean waveforms ± standard deviation for Clusters 1 and 3, showing distinct waveform morphologies with clearly different temporal profiles and amplitudes. **A(bottom)**. Inter-spike interval (ISI) distributions with kernel density estimation (KDE) for both clusters, demonstrating appropriate refractory period characteristics that confirm biological validity. **B(top)**. Waveform density plot for Cluster 1, revealing temporal stability and amplitude consistency across the recording session. **B(bottom)**. Waveform density plot for Cluster 3, providing complementary evidence for the distinct nature of this neuron through its characteristic waveform signature. **C(top)**. Eight exemplars that elicited a response (denoted by stars). This neuron responds to most pictures presented in the session, as suggested by the response pattern observed 500ms apart. **C(bottom)**. In contrast, the nearby neuron responded to a few pictures, including places, animals, and faces (4 non-responsive exemplars are also shown, denoted by circles). **D(top)**. Firing rate (Z-scored with respect to the baseline) for 228 stimuli shown in the session. 79.9% passed the response criterion, suggesting extremely low selectivity. The distribution of the Z-scores is shown in gold, with only 3% having Z < 5. Although ∼20% of the stimuli did not pass the criterion, the lack of separation between those and the responsive ones suggests the former are likely false negatives due to statistical fluctuations. **D(bottom)**. Only 2.2% (5 out of 228) of the stimuli passed the response criterion, and the separation in the distributions suggests no more than 4 or 5 potential false negatives. This high selectivity is also non-categorical. The presented non-responsive stimuli in **C(bottom)** are good representatives, as seen by the probability distribution.

## 3. Conclusion

In this work, we introduced **MCWs (MiCroWire sorter)**, a modular, open-source, and human-optimized spike sorting pipeline tailored for intracerebral microwire recordings. The framework is engineered to address the unique challenges of human electrophysiology, including sparse electrode arrays, dynamic environmental noise, and task-aligned data segmentation in clinical settings. Through its suite of configurable functional modules, spanning Data-Driven Noise Removal and Artifact Rejection to customizable Clustering and rigorous Quality Validation, MCWs presents a comprehensive pipeline to facilitate the accurate isolation of single-neuron activity even under challenging recording conditions. By integrating both established and novel methodologies in signal processing, clustering, and quality assessment, our pipeline ensures adaptability across diverse research paradigms and computational platforms. Notably, the pipeline’s compatibility with both Python and MATLAB allows broad accessibility, while its modular structure enables seamless integration of and customization for user-defined applications. This framework is a significant step toward closing the gap between non-human and human spike sorting approaches and the gaps in existing spike sorting algorithms. By enabling reproducible, high-quality single-neuron analyses from complex human data, MCWs empower users to explore the neural basis of cognition with greater precision and confidence.

## Software and Data Availability

The MCWs spike sorting pipelineis open-source and available on GitHub at https://github.com/Reylab/MCWs.

